# Conductance-based Adaptive Exponential integrate-and-fire model

**DOI:** 10.1101/842823

**Authors:** Tomasz Górski, Damien Depannemaecker, Alain Destexhe

## Abstract

The intrinsic electrophysiological properties of single neurons can be described by a broad spectrum of models, from the most realistic Hodgkin-Huxley type models with numerous detailed mechanisms to the phenomenological models. The Adaptive Exponential integrate-and-fire (AdEx) model has emerged as a convenient “middle-ground” model. With a low computational cost, but keeping biophysical interpretation of the parameters, it has been extensively used for simulations of large neural networks. However, because of its current-based adaptation, it can generate unrealistic behaviors. We show the limitations of the AdEx model and to avoid them, we introduce the Conductance-based Adaptive Exponential integrate-and-fire model (CAdEx). We give an analysis of the dynamics of the CAdEx model and show the variety of firing patterns it can produce. We propose the CAdEx model as a richer alternative to perform network simulations with simplified models reproducing neuronal intrinsic properties.

## 1 Introduction

Computational modeling of large scale networks requires to compromise between biophysical realism and computational cost. This requirement might be satisfied by many two-variable models (Morris & Lecar, 1981; Krinskii & Kokoz, 1973; Fitzhugh, 1961; E. M. Izhikevich, 2003; Brette & Gerstner, 2005). In particular, two-variable models that are largely used and studied are the Izhikevich model (E. M. Izhikevich, 2003) and the Adaptive Exponential Integrate and Fire (AdEx) model (Brette & Gerstner, 2005; Naud, Marcille, Clopath, & Gerstner, 2008; Touboul & Brette, 2008). The first variable of these models corresponds to the membrane voltage, the second, usually with slower kinetics, corresponds to neural adaptation and allows achieving more complex dynamics and firing patterns, which can be observed in neural recordings. In both models, the second variable has a form of an additional current flowing into the neuron, which may lead to unrealistic changes of membrane voltage, especially in the case of long and intense neuronal firings such as that observed during seizures (McCormick & Contreras, 2001). Here, we describe this limitation, and to avoid it, we propose a modification of the second variable dynamics. The adaptation variable in our model has the form of a conductance, introducing the *Conductance-based Adaptive Exponential Integrate and Fire* model or *CAdEx* model. Previous work has used conductance-based adaptation, but these models used were either more simplified (Treves, 1993) or much more detailed Hodgkin-Huxley type models with complex channel gating dynamics (Connor & Stevens, 1971). A universal phenomenological model linking f-I curves with adaptation dynamics (Benda & Herz, 2003) has a useful general form, but did not specify voltage dynamics and post-spike adaptation. The CAdEx model keeps the simplicity of the two-variables models, while extending the repertoire of possible subthreshold dynamics.

## 2 Conductance-based Adaptive Exponential model

The Adaptive Exponential Integrate and Fire model can reproduce many types of firing patterns: tonic spiking cells, regular spiking adaptive cells, bursting cells, etc (Naud et al., 2008; Clopath, Jolivet, Rauch, Lüscher, & Gerstner, 2007; Touboul & Brette, 2008). However, because the adaptation in AdEx model is a current, a strong and unrealistic hyperpolarization of the cell can appear after a period of prolonged firing (Fig. 1). Moreover, subthreshold adaptation in this model is linear, which means that in the periods of strong hyperpolarization the adaptation current does not deactivate and can remain unrealistically strong. For instance, modelling M-current mediated adaptation in this model poses a problem because M-channels are mostly closed when the membrane voltage remains below −60 mV (Brown & Adams, 1980). A similar problem happens for strong depolarization due to lack of saturation of the adaptation current. A corresponding problem exists for inward depolarizing currents such as the I_h_ current: HCN channels close above −70 mV and saturate below −80 mV (Banks, Pearce, & Smith, 1993).

**Figure 1:**
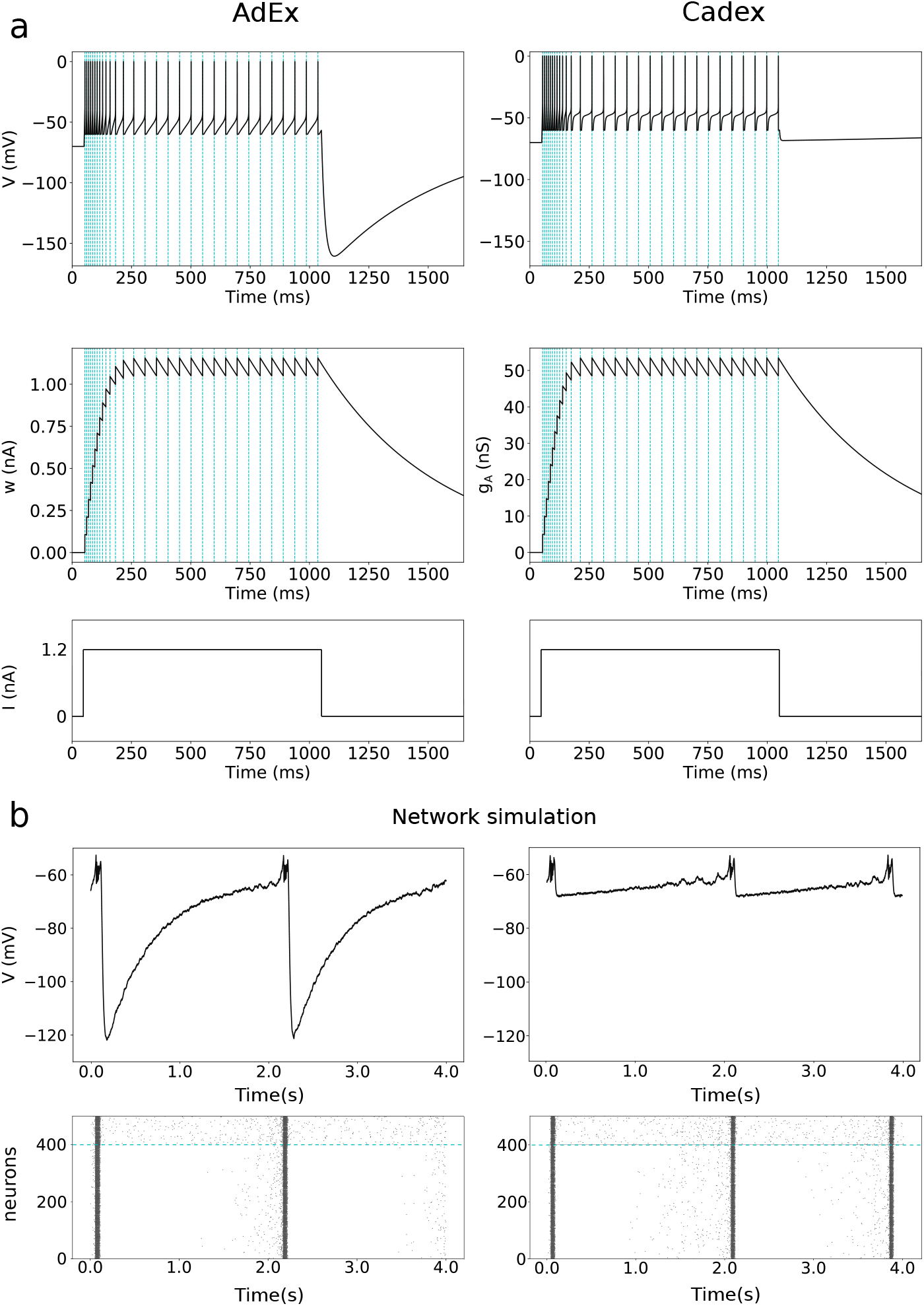
Comparison between AdEx and CAdEx models. (a) Both cells were injected with the same current pulse. Their firing rates *f* and adaptation parameters *A* were the same during pulse, *f* = 30 Hz and *A* = 0.03. However after pulse AdEx cell hyperpolarizes below −150 mV while CAdEx cell hyporpolarizes to proximity of the adaptation reversal potential, *E_A_* = −70 mV. The mathematical definition of the adaptation parameter, *A*, is given in Section 7. (b) Network simulations. The network comprises 1000 cells with 80% excitatory neurons with the same parameters values as for the corresponding single cell simulations. More details about these networks are given in Appendix A5. *Upper plots:* Mean membrane voltage of excitatory neurons. *Below*: Corresponding raster plots for 400 excitatory neurons (below blue line) and 100 inhibitory neurons (above blue line).

To overcome these problems, we propose here a model with a conductance based adaptation *g_A_* and with a sigmoid dependence of subthreshold adaptation on voltage. We named it Conductance based Adaptive Exponential model *CAdEx*. The equations of the model are as follows:

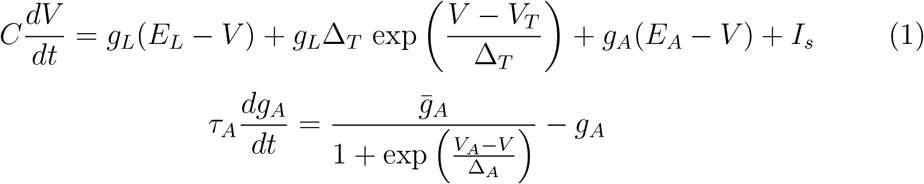

with after-spike reset mechanism:

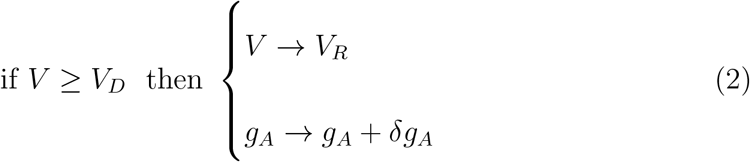

where *C* is the membrane capacitance, *g_L_* and *E_L_* are the leak conductance and reversal potential. *V_T_* is the spike threshold and Δ_*T*_ is the slope of the spike initiation, *E_A_* is the reversal potential of the adaptation conductance and *I_s_* is an input current. In the second equation, *τ_A_* is the time constant of adaptation, 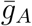 is the maximal subthreshold adaptation conductance, *V_A_* is the subthreshold adaptation activation voltage, and Δ_*A*_ is the slope of subthreshold adaptation.

A spike is initiated when *V* approaches *V_T_* and the exponential term escalates. After reaching a detection limit *V_D_*, the voltage is reset to the reset potential *V_R_* and it remains at this value during a refractory period Δ*t_ref_*. After each spike, *g_A_* is incremented by a quantal conductance *δg_A_*.

The sigmoidal subthreshold adaptation function is always positive, i.e. 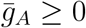, and it can be monotonically increasing, Δ_*A*_ > 0, or decreasing, Δ_*A*_ < 0, function of membrane voltage. In the simulations, the value of the detection limit has been set to *V_D_* = −40 mV and the value of the refractory period to Δ*t_ref_* = 5 ms.

In the Appendix A4, we exemplify the parametrizations for I_M_ and I_h_ currents based on biophysical data (Brown & Adams, 1980; Banks et al., 1993).

The already mentioned problem of unrealistic hyperpolarization in the AdEx model does not appear in the CAdEx model (Fig. 1). The corresponding problem of unrealistic hyperpolarization can appear also at the network level for AdEx neurons (Fig. 1b). Although both models can be tuned to exhibit very similar raster plots at the network level, the hyperpolarization problem disappears in networks of CAdEx neurons (Fig. 1). For a negative adaptation parameter *a* (corresponding to the accelerating firing pattern (Brette & Gerstner, 2005)) in the AdEx model, an additional problem of infinite hyperpolarization can appear, as shown in Appendix A6. Thus, the CAdEx model may be advantageous when the phenomena considered depend on the membrane potential.

All simulations of the models were done in Brian2 neural simulator and Python 3 programming language. The code which allows to run the CADEX model simulations can be downloaded from: https://github.com/neural-decoder/cadex.

## 3 Dynamical analysis of the model

### 3.1 Fixed points and bifurcations

The CAdEx is a nonlinear dynamical system with a discontinuous post-spike reset mechanism. The *V* and *g_A_* nullclines are as follows (Fig. 2):

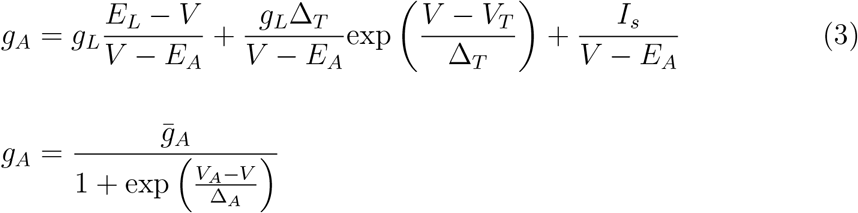

**Figure 2:**
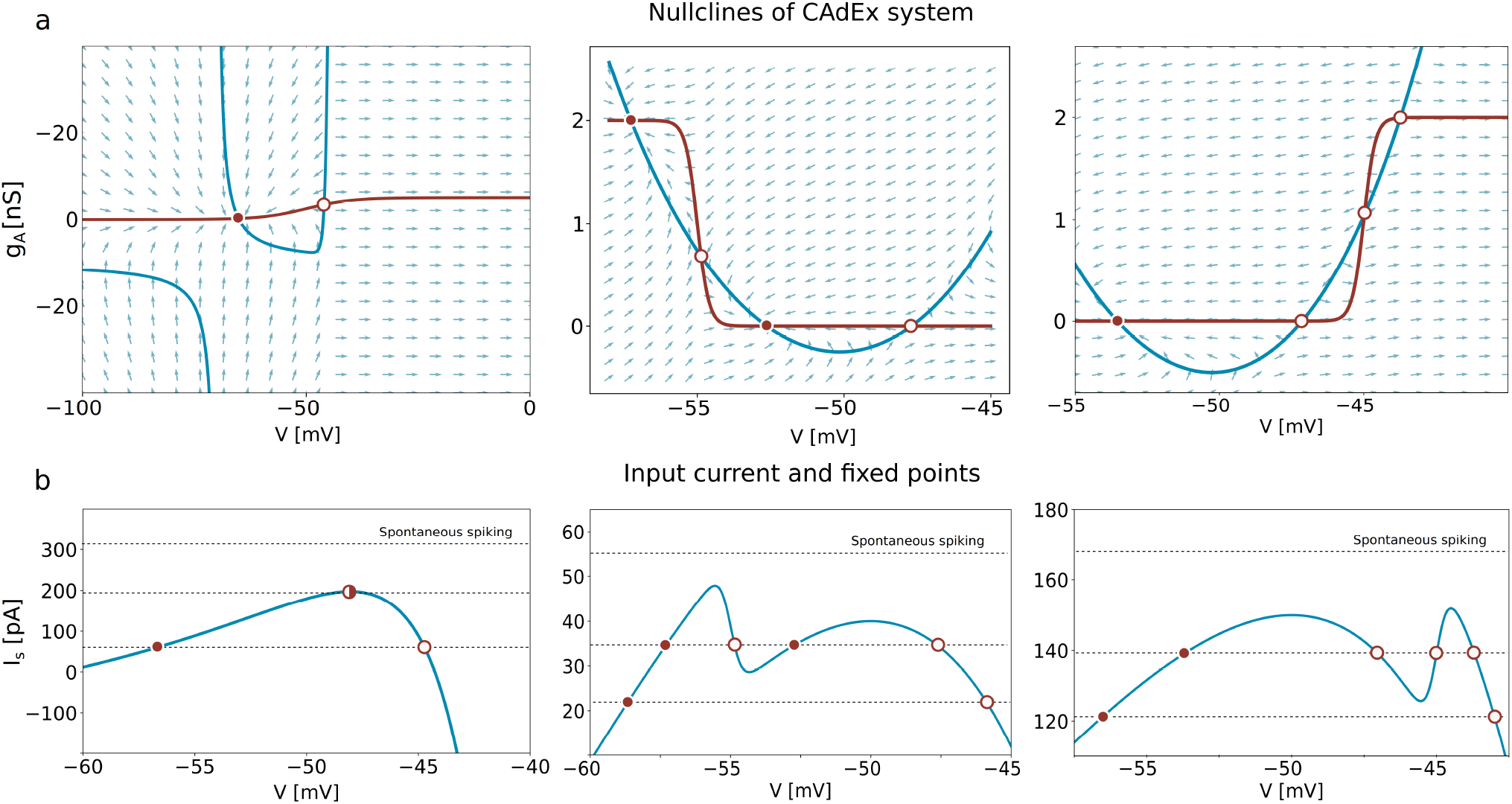
(a) Nullclines of the CAdEx system for three parametrizations, *left* for Δ_*A*_ > 0 and at most two fixed points, *center* for Δ_*A*_ < 0 and at most four fixed points, *right* for Δ_*A*_ > 0 and at most four fixed points. The blue line is a V-nullcline, red line-*g_A_*-nullcline. Red filled circle indicate stable fixed points, white filled circles indicate unstable fixed points, and, half-filled circles indicate neutral points. Arrows indicate the direction of the vector field. (b) Corresponding *S*(*V*) functions. Intersections with lines of constant input current *I_s_* indicate the positions of the fixed points.

The shape and the location of the *V*-nullcline depend on the input current *I_s_* (see Fig. A1 in Appendix A1). The *V*-nullcline is divided by the vertical asymptote *V* = *E_A_* into the left and right branches (see Fig. 2a.)

The intersections of nullclines give the fixed points of the system. From Eq. 3, the solutions of the equation

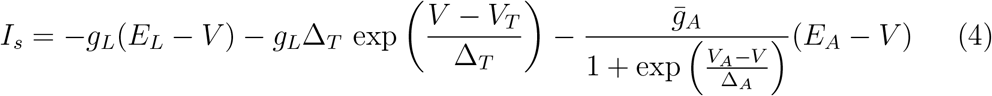

give the membrane voltages of the fixed points.

The analysis of the above equation, which we can write in a simpler form defining a new function *S*(*V*) as *I_s_* = *S*(*V*), gives us the number of fixed points for a given input current Fig. 2. The function *S*(*V*) tends to −∞ for *V* → ±∞ and it can have:

1. A single maximum *V_max_*, which corresponds to the configuration with at most two possible intersections between nullclines. Let *I_SN_* = *S*(*V_max_*). For a current input *I_s_* < *I_SN_* there are therefore 2 fixed points *V*^−^(*I_s_*) and *V*^+^ (*I_s_*), for *I_s_* = *I_SN_* there is one fixed point and for *I_s_* > *I_SN_* there is no fixed point. *I_SN_* is therefore the current above which the system always spikes spontaneously. The system can start to spike spontaneously for *I_s_* < *I_SN_*, if all fixed points lose stability before reaching *I_SN_* in an Andronov-Hopf bifurcation (see below).
2. Two local maxima 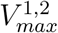, and one local minimum *V_min_*, which corresponds to maximally four possible fixed points. For input current *I_s_* < *S*(*V_min_*) there are two fixed points, one stable and one unstable. For 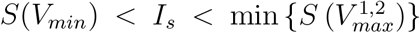 there are four fixed points, from which two can be stable. For 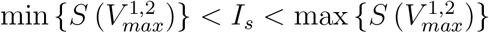 there are two fixed points, and for 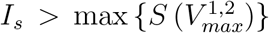 there is no fixed point and the neuron fires spontaneously.

To analyze the bifurcations of the system we need to consider the behavior of the Jacobian matrix at the fixed points *i* as a function of the input current, *J_i_*(*I_s_*) (see Appendix A2).

The trajectories in the phase space of trace 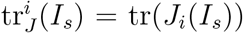 and determinant 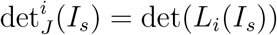 of the Jacobian give us the type of bifurcation, Fig. 3, 4.

In case (1), i.e. when there are maximally two possible fixed points, the value of tr_*J*_ at their merging point

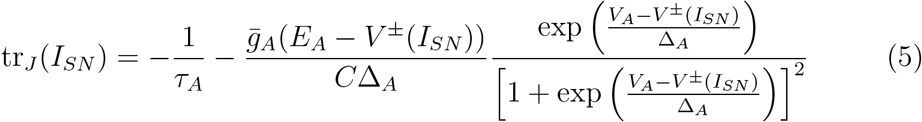

determines the type of bifurcation (see Appendix A2).

**Figure 3:**
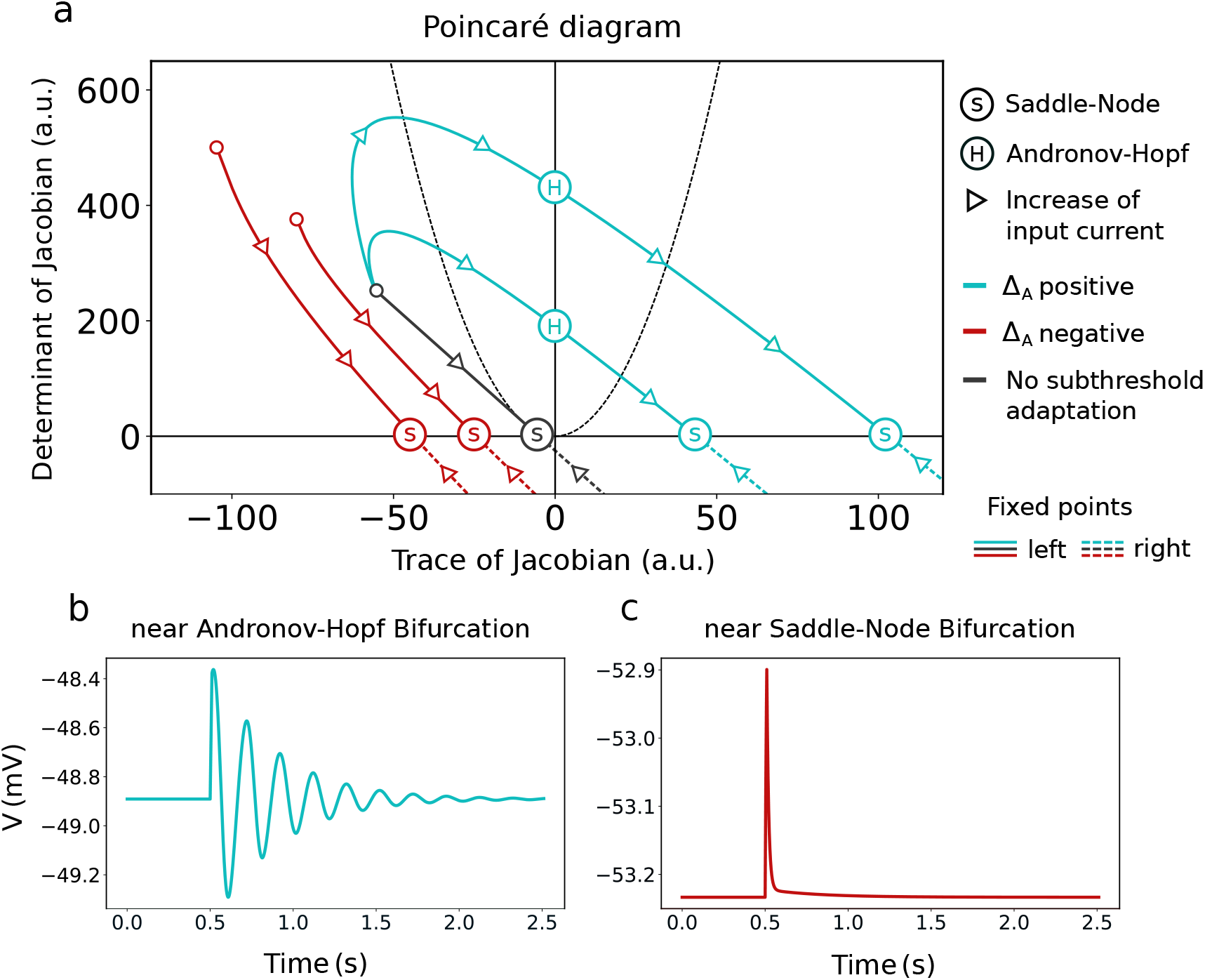
Bifurcations of the CAdEx system with at most 2 fixed points. (a) Trajectories of the fixed points on the Poincaré diagram for increasing input current *I_s_*. Solid lines denote trajectories of left, initially stable, fixed point with lower membrane voltage, dashed lines - trajectories of right, unstable fixed point with higher membrane voltage (shown partially). For positive Δ_*A*_, *blue lines*, the stable fixed point loses stability with an Andronov-Hopf bifurcation, 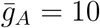 and 20 nS. For negative Δ_*A*_, *red lines*, the stable fixed point merges with the saddle point in a Saddle-Node bifuraction, 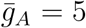 and 10 nS. Without subthreshold adaptation, 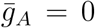 nS, *black line*, the system undergoes a Saddle-Node bifurcation. (b, c) Response of the cell to a brief current pulse, (b) near the Andronov-Hopf bifurcation, (c) near the Saddle-Node bifurcation. The parabola, *dashed black line*, is a function 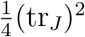, inside the area limited by the parabola the system can oscillate around equilibria.

If tr_*J*_(*I_SN_*) < 0, then the bifurcation is of Saddle-Node type. If tr_*J*_(*I_SN_*) = 0, the bifurcation is of Bogdanov-Takens type. If tr_*J*_(*I_SN_*) > 0, the bifurcation is of Andronov-Hopf type.

Since the merging point can be located only on the right branch of the V-nullcline, *E_A_* < *V*(*I_SN_*), then if Δ_*A*_ < 0, the bifurcation is always of Saddle-Node type. If Δ_*A*_ > 0, all types are possible depending on the sign of tr_*J*_(*I_SN_*). If 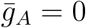, the bifurcation is also always of Saddle-Node type.

In case (2), i.e. when there are maximally four possible fixed points, two fixed points are created in a Blue Sky bifurcation when the input current reaches a minimum of *S*(*V*) function, *I_BS_* = *S*(*V_min_*), see Fig. 4. A Blue Sky bifurcation is a Saddle-Node bifurcation for an increasing input current in which two fixed points appear. If at the creation point tr_*J*_(*I_BS_*) < 0, a pair of stable and unstable fixed points are created. If tr_*J*_(*I_BS_*) ≥ 0, a pair of unstable fixed points are created. tr_*J*_(*I_BS_*) has the form:

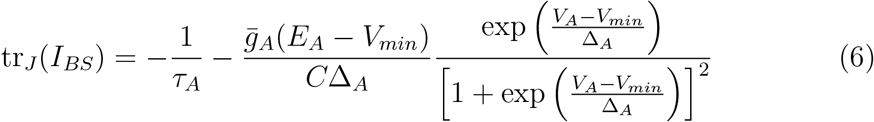

**Figure 4:**
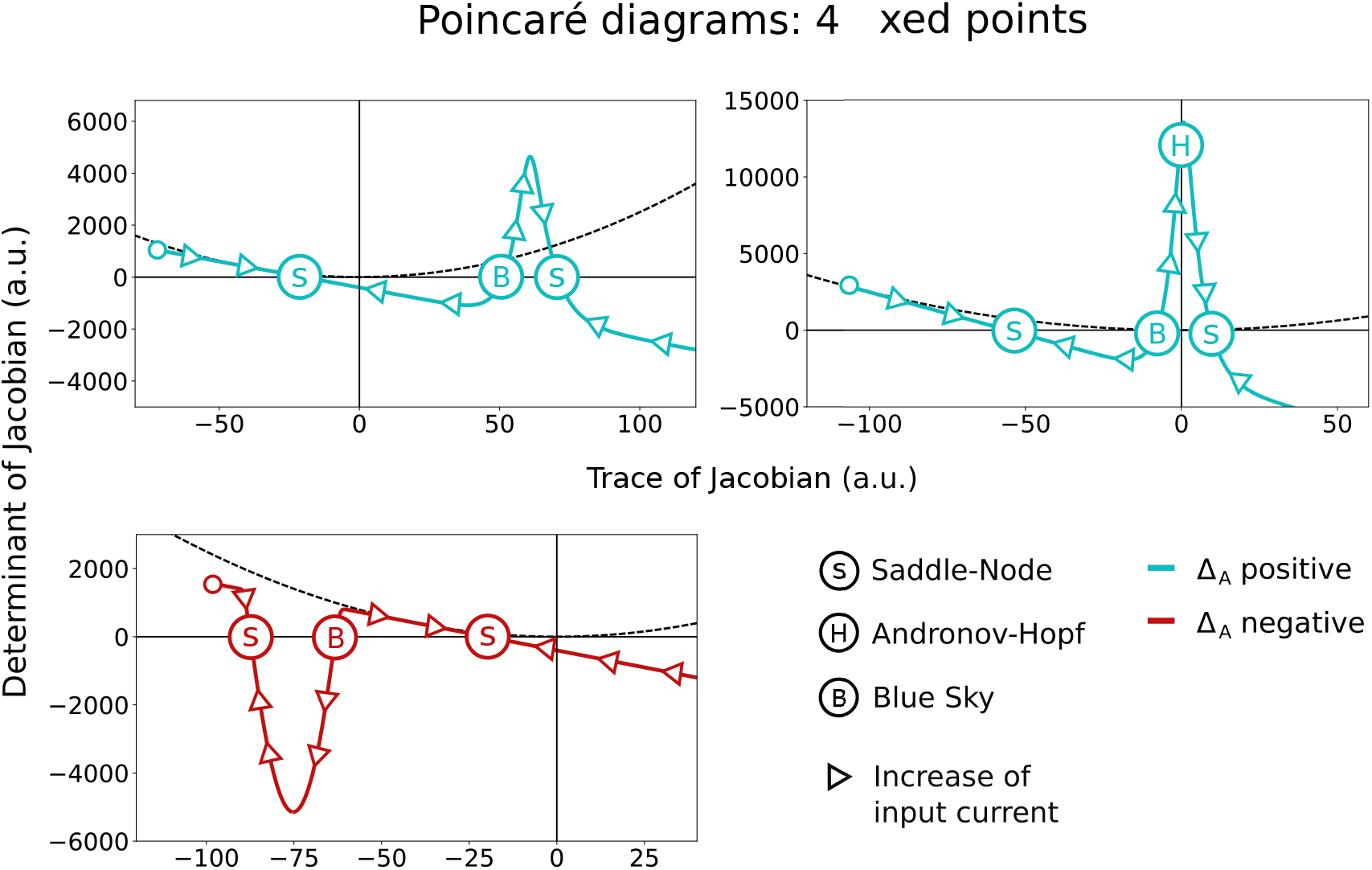
Bifurcations of the CAdEx system with at most 4 fixed points. Trajectories of the fixed points on the Poincaré diagram are shown for increasing input current *I_s_*. Trajectories are shown for positive Δ_*A*_, *blue lines*, for negative Δ_*A*_, *red lines*. Two fixed points are created in a Blue Sky bifurcation which subsequently merge with a stable and an unstable fixed point in a Saddle-Node bifurcation. *Upper right* plot: The stable fixed point created in a Blue-Sky bifurcation can lose stability in an Andronov-Hopf bifurcation before merging with the unstable fixed point in a Saddle-Node bifurcation. The parabola, *dashed black line*, is a function 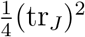, inside the area limited by the parabola the system can oscillate around equilibria.

If Δ_*A*_ < 0, the Blue Sky bifurcation point can be located only on the right branch of the V-nullcline and consequently *E_A_* < *V*(*I_BS_*) (center plot in Fig. 2). From Eq. 6 tr_*J*_(*I_BS_*) < 0 hence a stable and an unstable fixed points are created. If Δ_*A*_ > 0 the Blue Sky bifurcation point can be located on (1) the left branch of the V-nullcline for a strongly hyperpolarized neuron, *V*(*I_BS_*) < *E_A_* (Fig. A1 in Appendix A1). In this case, tr_*J*_(*I_BS_*) < 0 and a stable fixed point can be created. (2) the right branch of the V-nullcline, *V*(*I_BS_*) > *E_A_* (right plot in Fig. 4). In this case, both a pair of unstable fixed points or a pair of a stable and an unstable fixed point, can be created depending on the sign of tr_*J*_(*I_BS_*). If a stable fixed point is created it can subsequently lose its stability in an Andronov-Hopf bifurcation, see upper right plot in Fig. 4.

The trajectories of the equilibria and the corresponding dynamics in CAdEx model are much different than in the AdEx model. In the AdEx model at most only two fixed points are possible and the trajectories of equilibria on the Poincaré diagram are linear (see Appendix A.7).

### 3.2 Subthreshold oscillations

The system can oscillate around equilibrium if (a) the equilibrium is stable, i.e. tr_*J*_(*I_s_*) < 0 and det_*J*_(*I_s_*) > 0 and (b) eigenvalues of the Jacobian at fixed point have imaginary part, i.e. 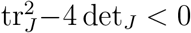. In that case, the frequency of oscillations is given by 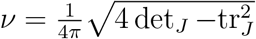, see Fig. 5 and Appendix A2.

**Figure 5:**
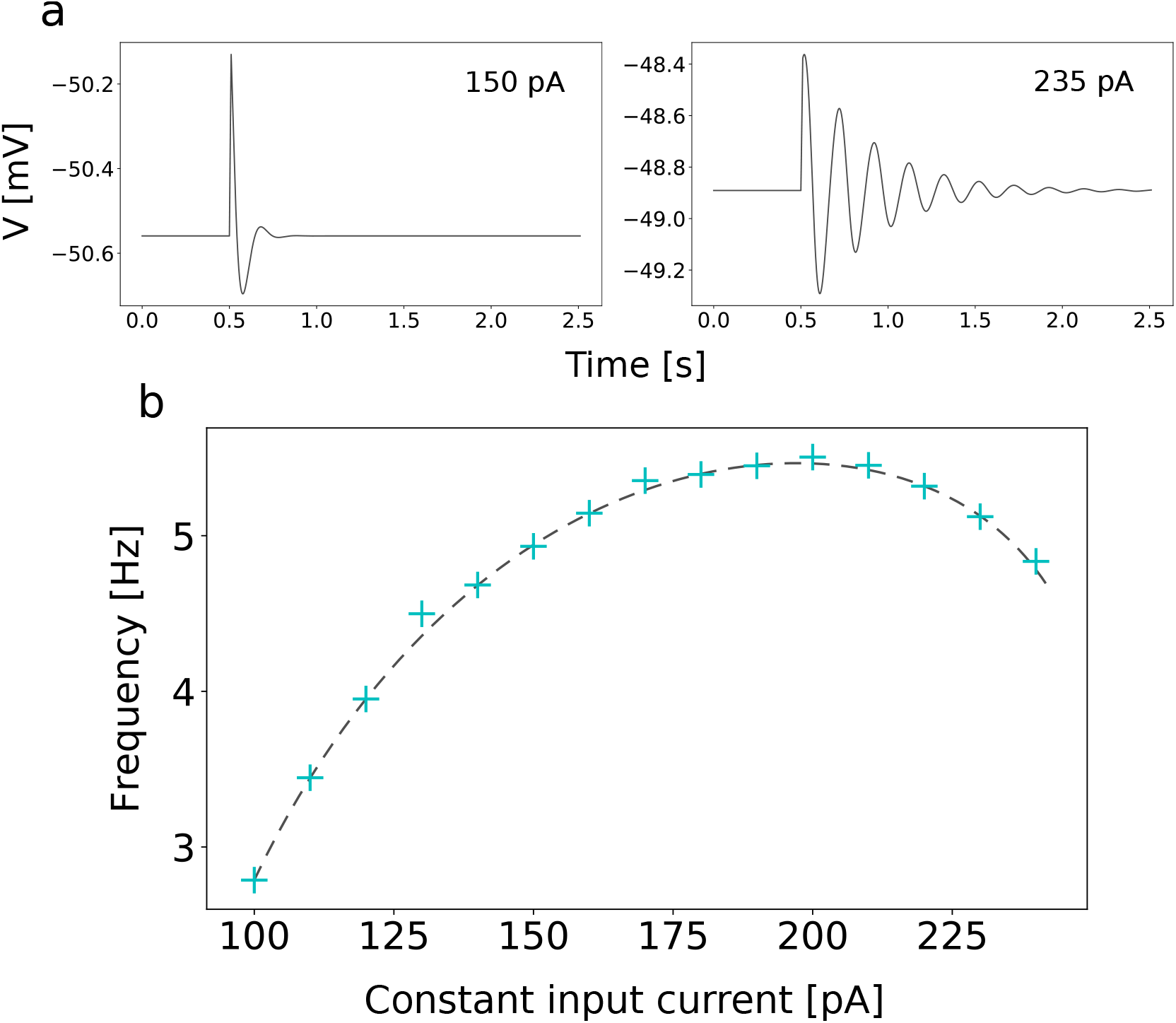
(a) Subthreshold oscillations of the membrane voltage around the stable equilibrium for different constant input currents. (b) Frequency of subthreshold membrane voltage oscillations in the CAdEx model as a function of a constant input current. The crosses signify numerical results, a dashed line - theoretical prediction, 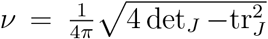. Here, a pulse of current of amplitude 10 pA and duration of 10 ms was applied when the membrane potential was at a stable equilibrium.

The sustained subthreshold oscillations due to the emergence of a limit cycle can be also observed. Some examples in the case of multi-stability are given in the next section.

## 4 Multi-stability

In this section, we present a list of examples of multi-stability. Such behaviors can be observed due to two properties of the model.

### Subthreshold multi-stability

Due to the non-linear form of the subthreshold adaptation, the system may exhibit two or four fixed points. With four fixed points, various configurations are possible: with a positive or negative slope (Δ_*A*_) of the adaptation nullcline. The stability of these fixed points, depending on the input current, are shown in Fig. 4.

### Superthreshold stability

The model has a discontinuity which occurs after a spike and corresponds to the reset of the membrane voltage and to the increment of adaptation conductance. It can lead to steady-state firing behavior, which can be treated as an attractor of the model. The reset discontinuity has, in many aspects, the same effect as a third variable. It allows chaotic behaviors (as described in the next section) normally impossible in a continuous two dimensional system.

Here, we give three examples of multi-stability: (a) with four fixed points and negative slope of adaptation, (b) with four fixed points and positive slope of adaptation and (c) with two fixed points and positive slope of adaptation.

a. With a negative and small enough value of Δ_*A*_, the nullclines can have four crossings. In this case, the system has two stable fixed points with different values of the membrane potential. In this situation, three stable steady-states are observed, Fig. 6a: Two possible resting states of membrane potential and a self-sustained spiking state. Small perturbations permit a switch between these steady-states, as shown in Fig. 6b. By applying synaptic noise through conductance-based inhibitory and excitatory synapses (see Appendix A3), transitions between these stable states can be observed, Fig. 6c.
b. With a positive and small enough value of Δ_*A*_, the system can also have four fixed points. In this case, the emergence of a stable limit cycle is observed. It leads to three possible stable behavior: resting state, subthreshold oscillations, and regular spiking as show in Fig. 7a. As in the previous condition, transitions between these states can be observed under synaptic bombardment, Fig. 7b.
c. In a situation with two fixed points, the emergence of an unstable limit cycle is observed, leading to bi-stability near the threshold between spiking and, non-spiking damped oscillations, as shown in Fig. 8c.

**Figure 6:**
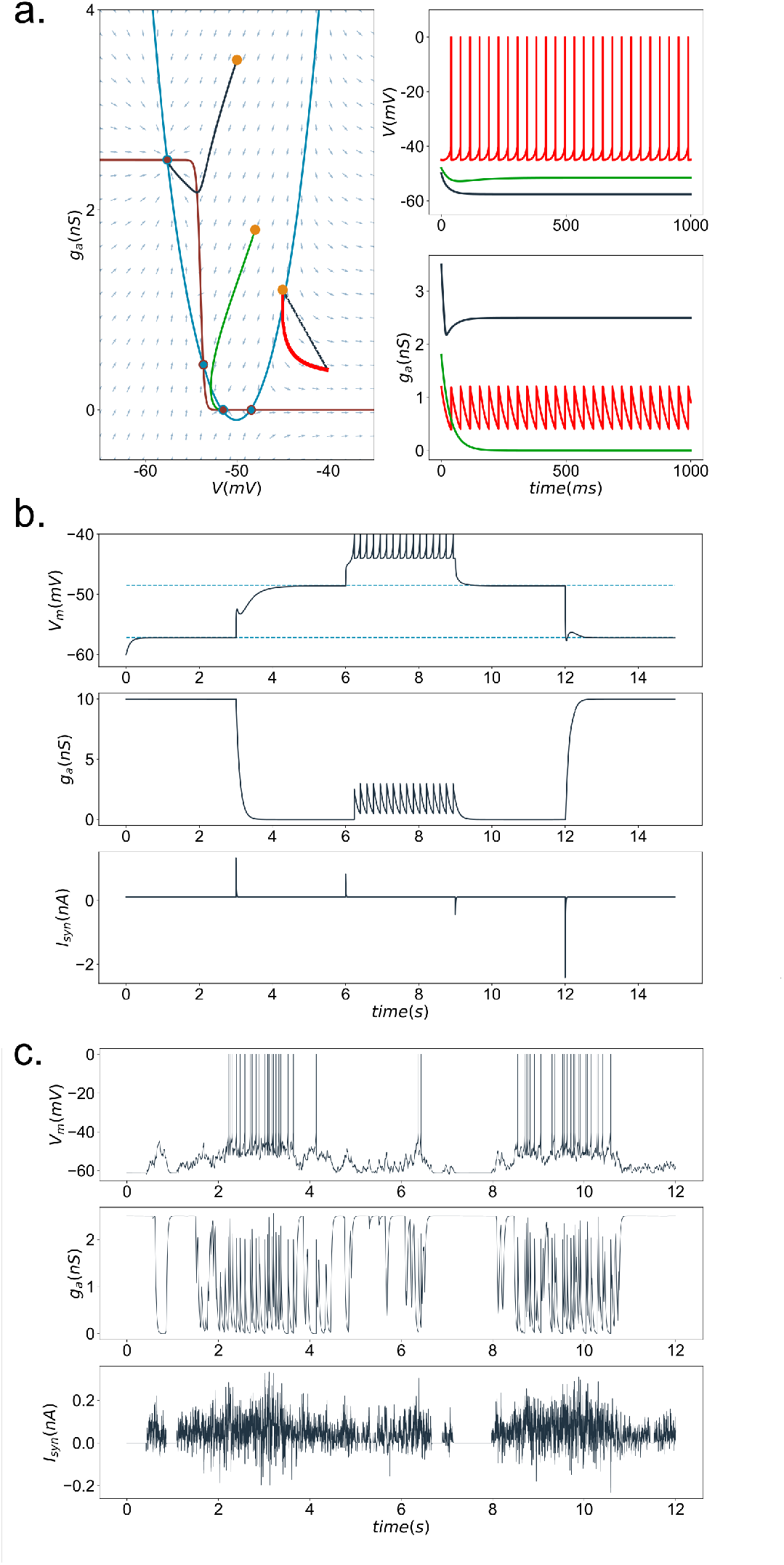
Multi-stability observed with a negative slope of *g_A_*: (a) the dynamics for single neuron with two stable fixed-points and a stable limit cycle due to the reset, (b) transitions between possible steady-states due to excitatory or inhibitory short-time perturbations, (c) transition between up and down state of single neuron receiving input through conductance-based synapses.

**Figure 7:**
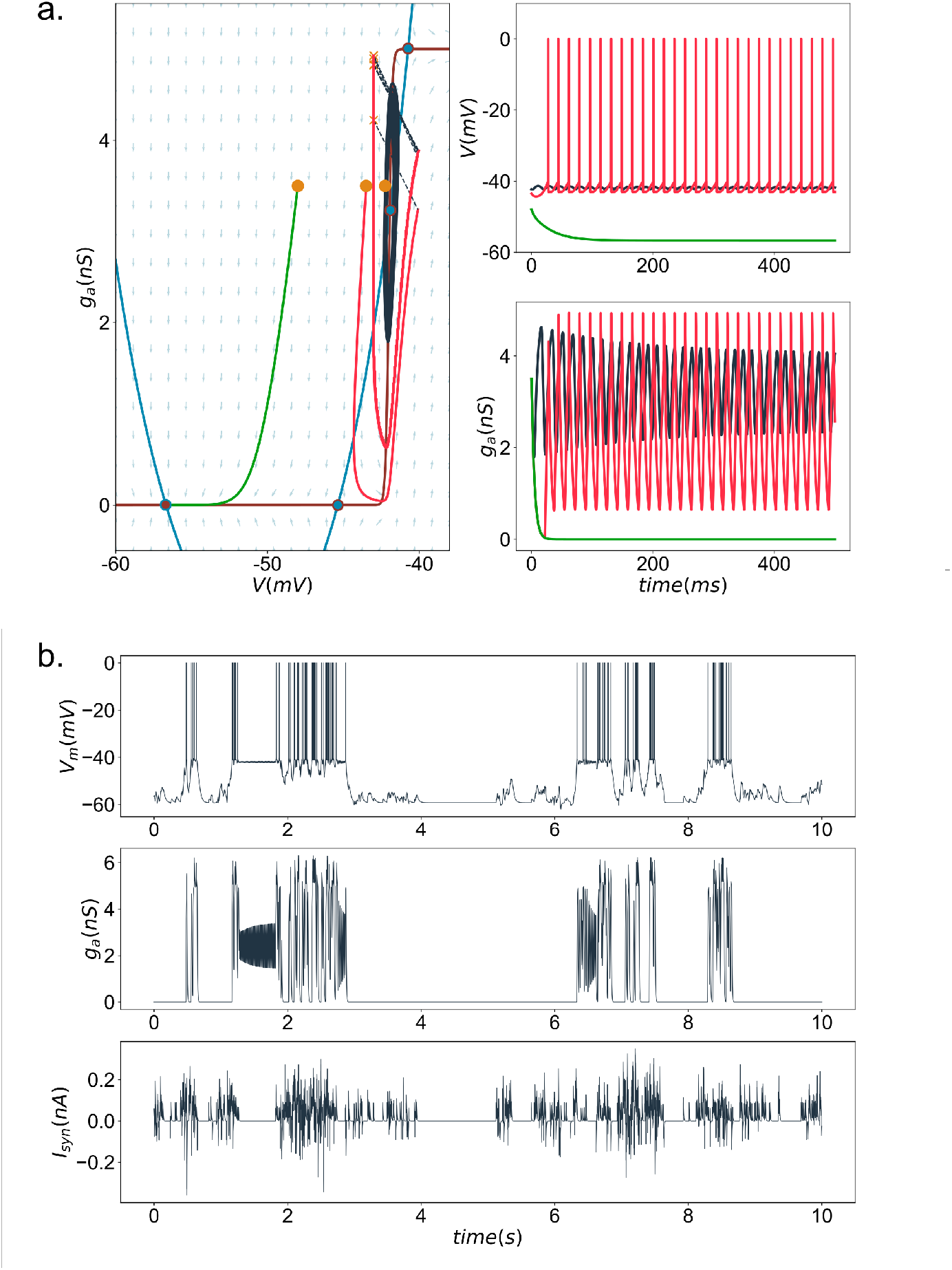
Multi-stability observed with a positive slope of *g_A_*: (a) the dynamics for single neuron with a stable fixed-point: a stable limit cycle due to the subthreshold dynamics and a limit cycle due to the reset, (b) transitions between up and down state of single neuron receiving input through conductance-based synapses.

**Figure 8:**
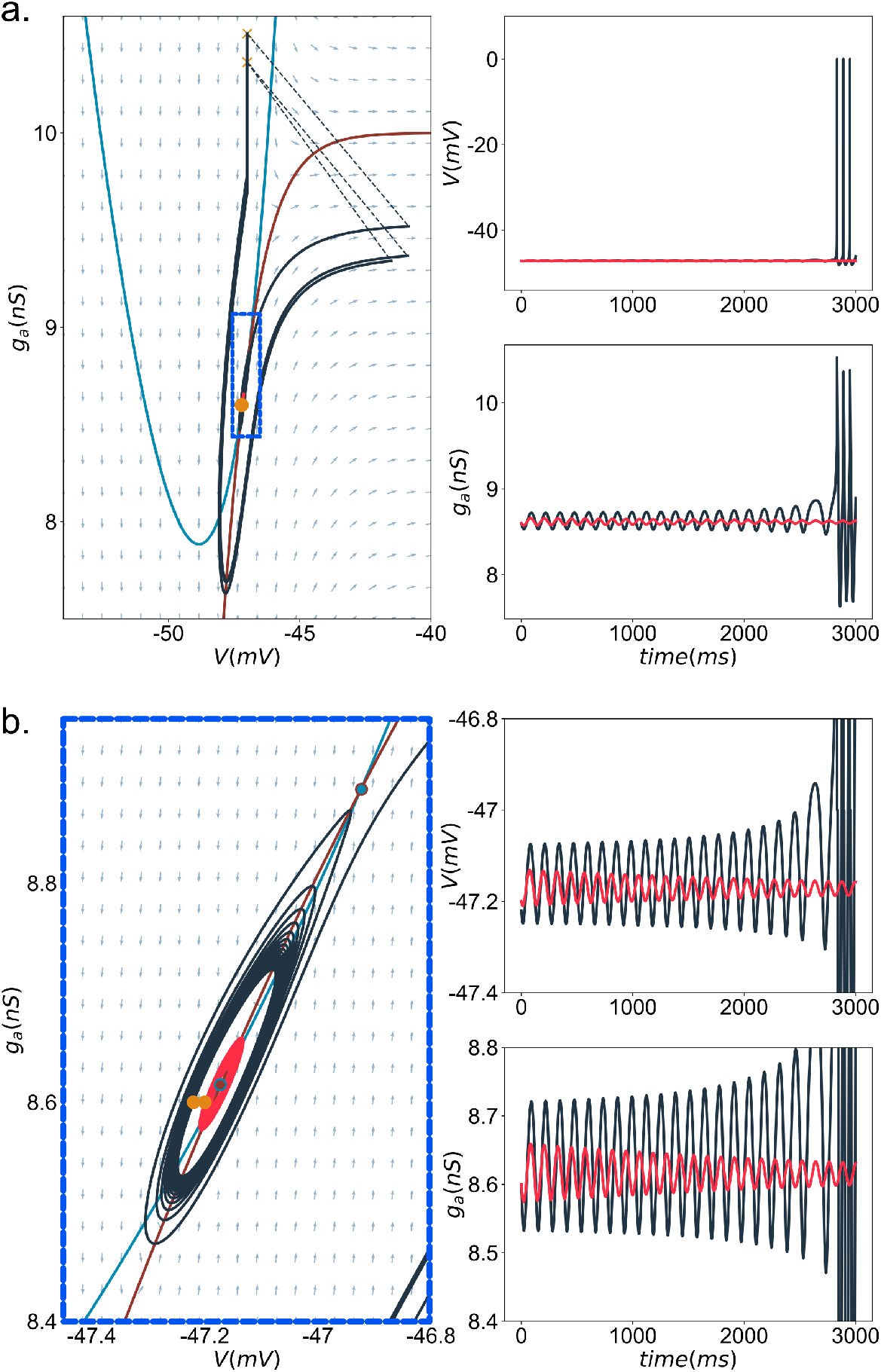
Multi-stability observed with two fixed points, (a) damped subthreshold oscillations are observed near the spiking behavior. (b) Enlarged concerned region (blue square). Two simulations initiated: inside (red) and outside (dark grey) the unstable limit cycle

## 5 Firing patterns

The CAdEx model is able to reproduce a large repertoire of intrinsic electrophysiological firing patterns. In this section, the dynamics of firing patterns in CAdEx, reproducing those found in AdEx, will be detailed. In order to facilitate the use of this model to reproduce specific behaviors, we describe which parameters influence specific firing patterns. The initial values of the two variables in the simulations were *V*(0) = −60 mV for the membrane potential and *g_A_*(0) = 0 nS for the adaptation; except when Δ_*A*_ < 0 when *g_A_* is initialized at 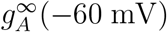. For the sets of parameters used in this section, the system does not have fixed points. Therefore, independently of the starting point location, the system converges to the same steady-state firing. However behaviors in the transient regime may change, which is crucial for spike frequency adaptation. The parameters for each pattern of Fig. 9 are given in Table 1.

**Figure 9:**
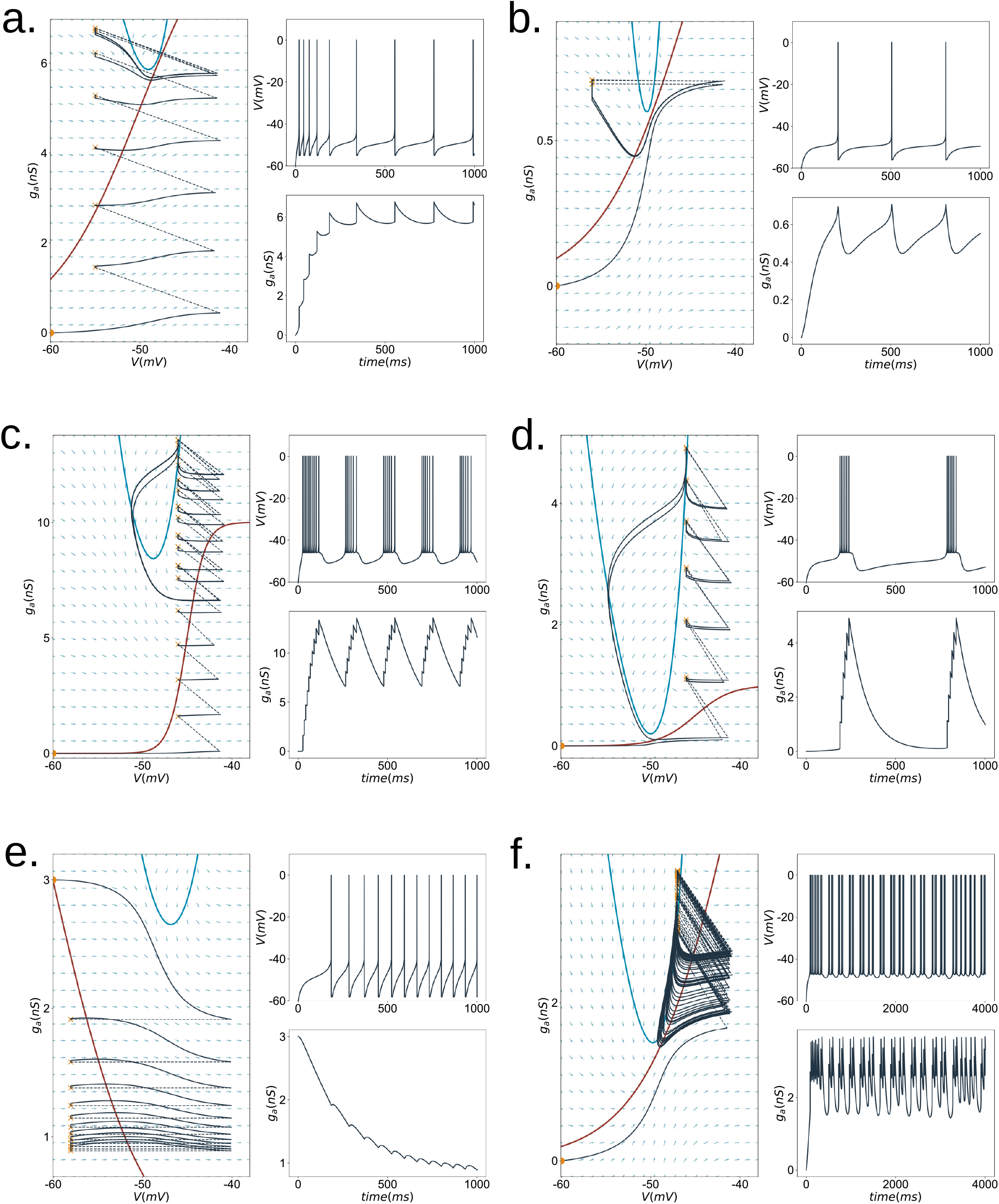
Firing patterns of the CAdEx model: (a) adaptive spiking, (b) delayed low-frequency tonic spiking, (c) bursting, (d) delayed bursting, (e) accelerated spiking, (f) chaotic-like spiking (see Table 1 for parameters).

**Table 1:** Parameters of the CAdEx model for the firing patterns shown in Fig. 9

| | $C_m$ [pF] | $E_A$ [mV] | $E_L$ [mV] | $I_s$ [pA] | $V_A$ [mV] | $\Delta_A$ [mV] |
| --- | --- | --- | --- | --- | --- | --- |
| Adaptive spiking | 200 | -70 | -60 | 200 | -50 | 5 |
| Tonic spiking | 200 | -70 | -70 | 192 | -45 | 5 |
| Bursting | 200 | -60 | -58 | 150 | -45 | 1 |
| Delayed bursting | 200 | -70 | -60 | 100 | -45 | 2 |
| Accelerated spiking | 200 | -70 | -60 | 130 | -60 | -5 |
| Chaotic-like spiking | 200 | -70 | -58 | 90 | -40 | 5 |

| | $V_R$ [mV] | $V_T$ [mV] | $\delta g_A$ [nS] | $\bar{g}_A$ [nS] | $g_L$ [nS] | $\tau_A$ [ms] |
| --- | --- | --- | --- | --- | --- | --- |
| Adaptive spiking | -55 | -50 | 1 | 10 | 10 | 200 |
| Tonic spiking | -56 | -50 | 0 | 2 | 10 | 40 |
| Bursting | -46 | -50 | 1 | 10 | 10 | 200 |
| Delayed bursting | -46 | -50 | 1 | 1 | 12 | 100 |
| Accelerated spiking | -58 | -48 | 0 | 6 | 10 | 300 |
| Chaotic-like spiking | -47 | -50 | 1 | 10 | 10 | 25 |

### Adaptive spiking

The adaptation can cause a decrease of spiking frequency as seen in Fig. 9a. Correspondingly the value of *g_A_* increases until it reaches a steady state. The spike frequency adaptation is mainly affected by the spike triggering parameters (*V_T_* and Δ_*T*_), by the membrane properties: *C* and *g_L_*, as well as by the adaptation parameters, 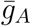 and strongly by 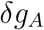.

### Tonic spiking

In the absence of adaptation, i.e. 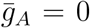, the model exhibits tonic spiking. However, tonic spiking can also be observed with non-zero adaptation, as in Fig. 9b, if the steady state firing can be reached after the first spike. The steady state spiking frequency is influenced by the membrane capacitance, *C*, the leak conductance, *g_L_*, and the slope of spike initiation Δ_*T*_, and also by the adaptation time constant *τ_A_*. The after-hyperpolarization following spiking depends on both: reset value *V_R_* and adaptation time constant *τ_A_*.

### Bursting

To obtain bursting behaviors, Fig. 9c, the reset value, *V_R_*, has to be higher than the voltage of the minimum of the *V*-nullcline. The system spikes until it crosses the *V*-nullcline, ending the burst. Then, above the *V*-nullcline, the system is driven to lower values of adaptation and membrane potential. After a second crossing of the *V*-nullcline, the system spikes again, starting a new bursting cycle. Bursts can be characterized by intra- and inter-bursts activities. The interburst activity is affected by adaptation, 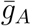 and *τ_A_* strongly determine the interburst time interval and after burst hyperpolarization.

### Delayed spiking

All firing patterns can occur with a time delay. Example is given for bursting, Fig. 9d, and low frequency spiking Fig. 9b. The distance between *V* and *g_A_* nullclines is determined by *I_s_*. By changing this distance and the time constant *τ_A_* it is possible to obtain a region of slow flow (small 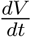) and consequently to increase the delay and the inter-event interval. The reset *V_R_* and *δg_A_* also affects the time required for the system to go around the *V* nullcline, and then, to the next event.

### Accelerated spiking

A negative slope of *g_A_* (Δ_*A*_ < 0) associated with *δg_A_* = 0, leads to firing rate acceleration as shown in Fig 9e.

### Chaotic-like spiking

As the CAdEx model is not continuous due to the after spike reset of voltage and incrementation of adaptation, chaotic-like spiking behavior can be observed as shown in Fig. 9f. These phenomena occur when the reset is close to the right branch of the *V* nullcline. To verify that this irregularity is not due to numerical error, we used various integration methods and various time steps in Brian2 simulator (Stimberg, Brette, & Goodman, 2019).

## 6 Adaptation and irregularity

A convenient way to measure spike frequency adaptation is given by the adaptation index:

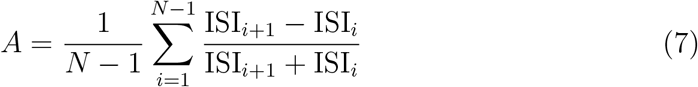

where *N* is the number of subsequent interspike intervals, ISI_*i*_. Note that *A* ∈ (−1, 1) and is positive for decelerating spike trains and negative for accelerating spike trains.

To measure the irregularity of spiking, we used the coefficient of variation of ISI:

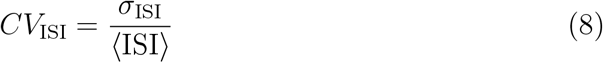

where 〈ISI〉 and *σ*_ISI_ are the mean and the standard deviation of ISIs respectively. According to this definition, *CV*_SI_ ∈ [0, ∞), where *CV*_SI_ = 0 for regular tonic spiking.

Both subthreshold 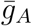 and post-spike *δg_A_* adaptation parameters affect the adaptation index, see Fig. 10a. This allows the model to reproduce the wide range of A index values observed in neurons (Allen Brain Institute, 2015).

**Figure 10:**
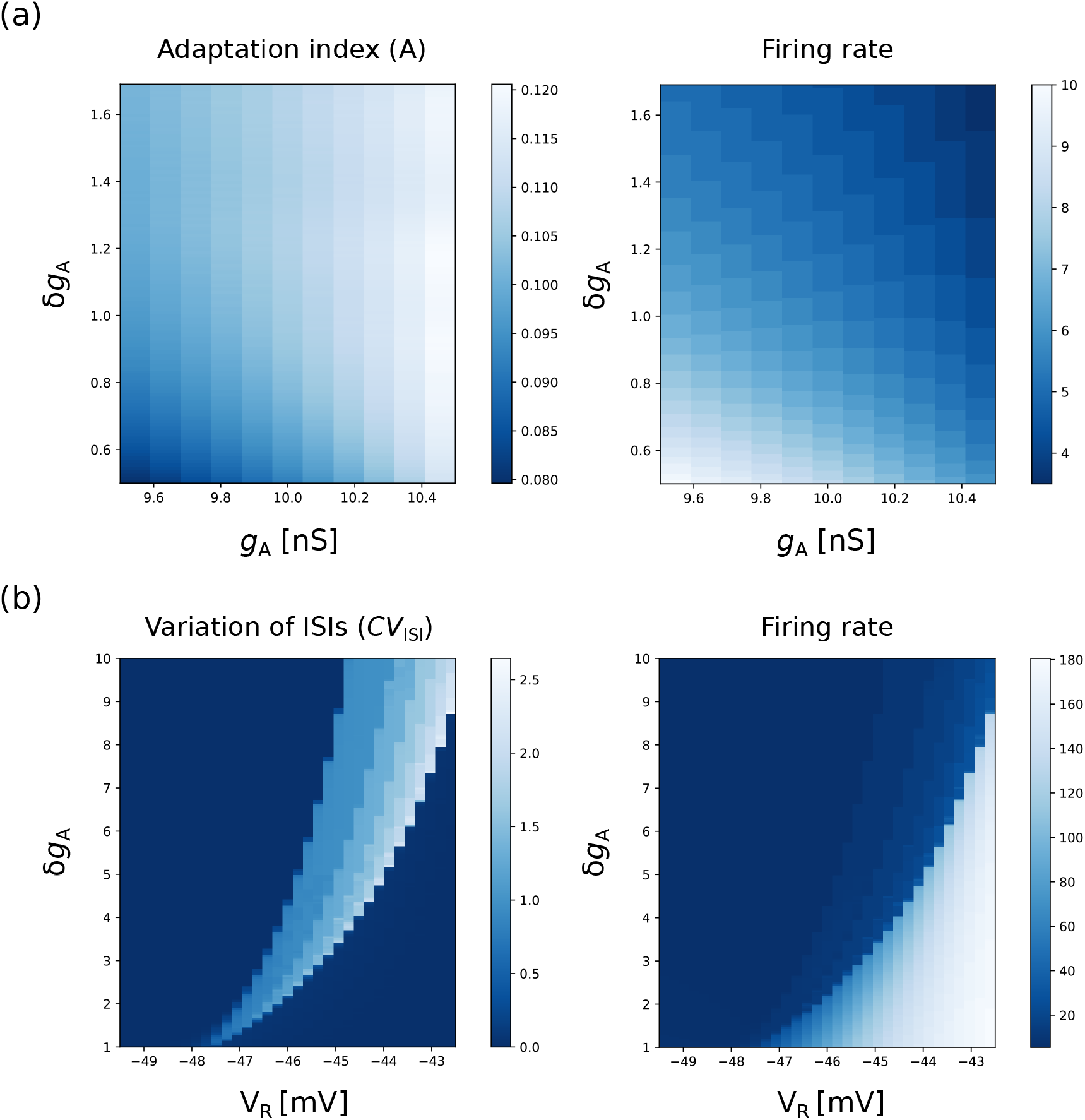
Spike frequency adaptation and variation of interspike intervals in the CAdEx model. (a) Adaptation index *A* and corresponding firing rate as a function of the maximal subthreshold adaptation 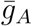 and post-spike adaptation *δg_A_*. (b) Coefficient of variation of interspike intervals *CV*_ISI_ and corresponding firing rate as a function of reset potential *V_R_* and post-spike adaptation *δg_A_*.

The irregular spiking (like chaotic-like spiking and bursting) is especially pronounced in the transition zone between slow and fast regular spiking regions, see Fig. 10b. On the phase diagram, slow regular spiking corresponds to a post-spike reset occurring on the left side of the right branch of the V-nullcline, and, consequently, leading to longer interspike intervals, while fast tonic spiking corresponds to a reset occurring on the right side, leading to fast subsequent spike. In the transition zone alternations between resets on the left and right sides of the V-nullcline can lead to highly irregular spiking (cf.chaotic-like Fig. 9f) and bursting Fig. 9c and d).

This irregularity of spiking originates from the post-spike reset mechanism and it also occurs in the AdEx model (Touboul & Brette, 2008). The correspondence between parameters of both models is intuitive: *δg_A_* corresponds to *b* in the AdEx model, while the reset potential *V_R_* has the same meaning in both models.

## 7 Discussion

In this paper, we have proposed a new integrate-and-fire model with two variables, and which can produce a large repertoire of electrophysiological patterns while still allowing for clear mathematical insights and for large scale simulations. This CAdEx model is completely specified with twelve biophysical parameters, and reproduces qualitatively similar pattern as the AdEx model (Naud et al., 2008), because the *g_A_* nullcline may be considered as locally linear and approximates that of the AdEx model. While the dynamics of the CAdEx model is comparable to the AdEx model for moderate input and firing, the CAdEx model does not suffer from an unnaturally strong hyperpolarization after prolonged periods of strong firing. This can be very advantageous for modelling of highly synchronized rhythms and firings, like slow-wave oscillations or epileptic seizures. The different behaviors of the averaged membrane voltages between the models was demonstrated here at the network level. Moreover, the sigmoidal subthreshold adaptation function allows one to model the dynamics of voltage dependent ion channels in more detail, while retaining the overall computational simplicity. The sigmoidal form of the adaptation function enriches the dynamics, allowing a wider repertoire of multi-stabilities. An important difference between CAdEx and AdEx models is that the CAdEx model includes a new bifurcation structure with an Andronov-Hopf bifurcation in the 4-fixed point configuration (see Fig. 4, upper right). In this regime, there are two simultaneously stable fixed points, one can be in an “integrator” mode (and so going through saddle-node bifurcation when loosing stability) with the other one in a “resonator” mode, and the strongly resonating fixed point can then undergo an Andronov-Hopf bifurcation (E. Izhikevich, 2007). Thus, the CAdEx model can display a bistability between integrator and resonator modes for the same set of parameters. It would be interesting to see if such a predicted bistability can be found in other models, or experimentally.

The CAdEx model also solves a problem of divergence of the AdEx model to minus infinity for a negative slope of subthreshold adaptation. This divergence problem can be a source of instabilities in network simulations. It is automatically fixed by the CAdEx model because the voltage is bounded to the reversal value of the adaptation current.

However, our model has some limitations. First, the adaptation has the form of a non-inactivating current (such as I_M_ potassium current) which limits the description of a class of inactivating ionic channels. It also includes only one type of subthreshold adaptation. In comparison to the AdEx model, the computational cost of our model may be slightly higher due to the form of the adaptation variable, and more specifically the introduction of an exponential function. Also, as more parameters are introduced, more thorough dynamical studies and explorations of the parameter space are needed.

## Acknowledgement

The work was supported by CNRS, the European Community (Human Brain Project, H2020-785907), and the ICODE excellence network.

## A Appendix

### A.1 Shape of V-nullclines while varying input current

The V-nullcline changes its shape when the input current *I_s_* changes. When the input current is strong and hyperpolarizing, the V-nullcline is inverted near an asymptote *V* = *E_A_*, see Fig. A1.

**Figure A1:**
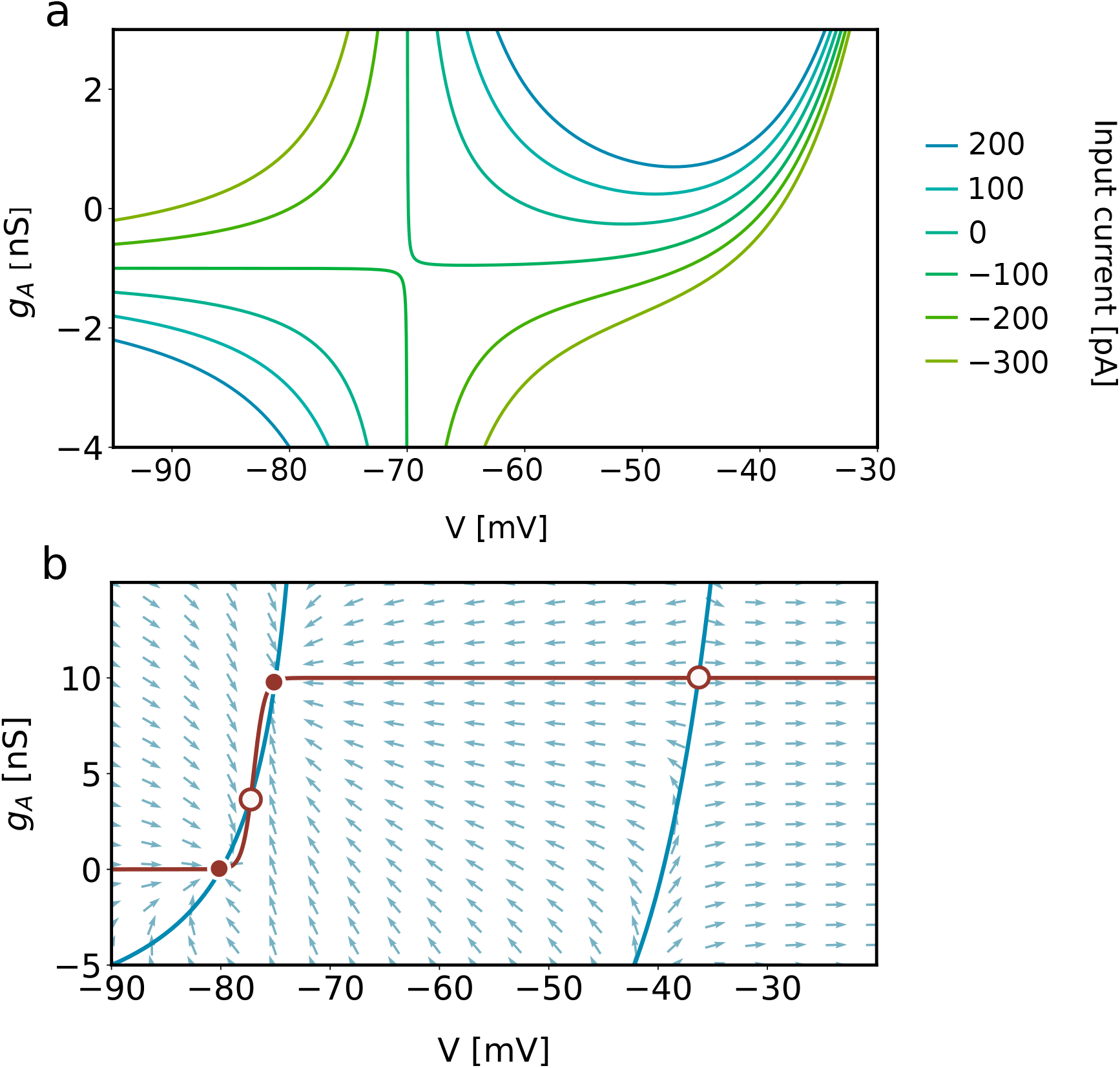
(a) The V-nullclines while varying input current *I_s_*. For strong and hyporpolarizing input current V-nullcline is inverted near an asymptote *V* = *E_A_* = −70 mV. (b) The four fixed points of the system after inversion of the V-nullcline by strong, hyperpolarizing current, *I_s_* = −200 mV

### A.2 Bifurcation analysis

#### Minima and maxima of *S*(*V*) function

The derivative of the function *S*(*V*) is given by

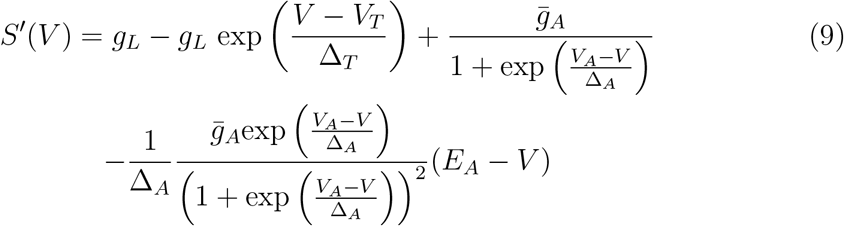

The solutions of the transcendental equation *S*′(*V*) = 0 give the locations of

1. a single global maximum of *S*(*V*) function, *S*(*V_max_*), for two possible intersections between the *V* and *g_A_* nullclines.
2. two maxima and one minimum of *S*(*V*) function, for four possible intersections between the *V* and *g_A_* nullclines.

#### Local linearized dynamics around equilibria

The Jacobian of the CAdEx system around equilibrium *i*, located at 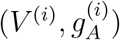, for an input current *I_s_* has the form:

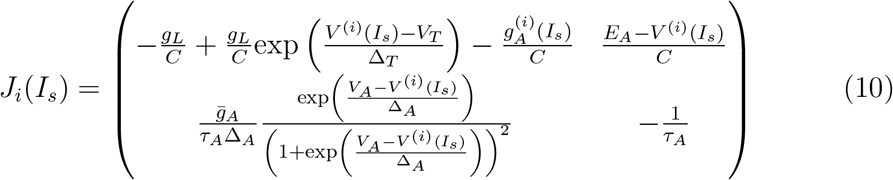

The trace 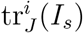 and the determinant 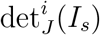 of the Jacobian are as follows:

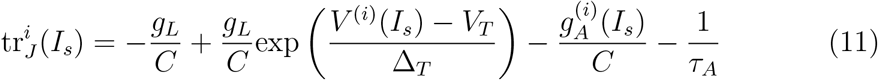

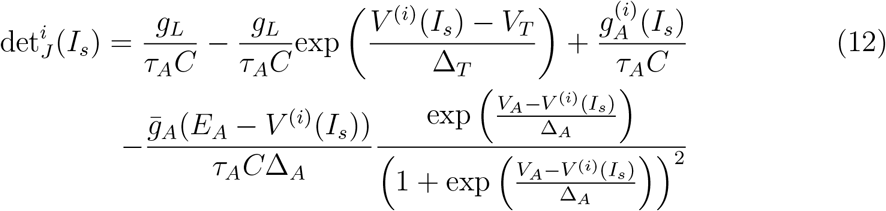

From the above equations, we obtain:

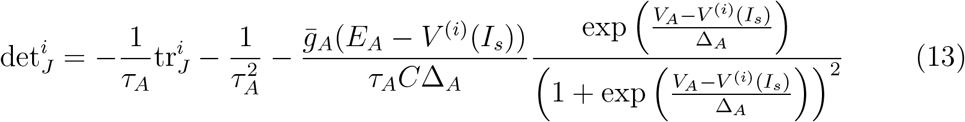

Let’s consider the case with at most two possible fixed points {−, +}. For *I_s_* → −∞, *V* of the left (−) fixed point tends to −∞. Since *g_A_* ≥ 0, from Eq. 10 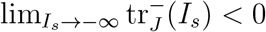. Similarly from Eq. 11 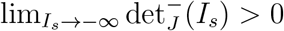. It means that the trajectory of the left fixed point on the Poincaré diagram 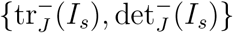 always starts in the stable quadrant (tr_*J*_ < 0, det_*J*_ > 0).

The trajectory of the left fixed point ends when it merges with an unstable fixed point on the axis det_*J*_ = 0. It happens for an input current *I_SN_*. If tr_*J*_(*I_SN_*) < 0 the bifurcation is of Saddle-Node type. If tr_*J*_(*I_SN_*) = 0, the bifurcation is of Bogdanov-Takens type. If at the merging point tr_*J*_(*I_SN_*) > 0, it means that the trajectory crossed the axis tr_*J*_ = 0 before, therefore the fixed point lost its stability in an Andronov-Hopf bifurcation. From Eq. 12

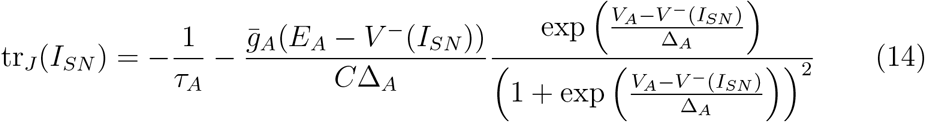

The eigenvalues of the Jacobian matrix at fixed points are then given by:

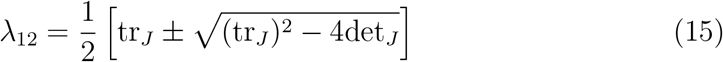

If the eigenvalues are complex, i.e. (tr_*J*_)^2^−4det_*J*_ < 0, then the system oscillates around equilibrium. The imaginary part of an eigenvalue is equal to the angular frequency of the oscillation, i.e. Im(λ) = *ω* = 2*πν*. Consequently the frequency of oscillations is given by:

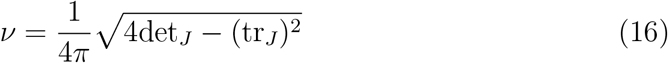

### A.3 Conductance-based synapses

To describe the behavior of the system receiving synaptic input, as in Section 4 concerning Multi-stability, we used a conductance-based model of synaptic inputs.

The synaptic input current to our model is given by the following equation:

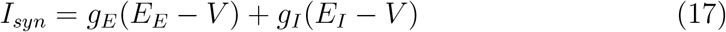

Where *E_E_* = 0 mV is the reversal potential of excitatory synapses and *E_I_* = −80 mV is the reversal potential of inhibitory synapses. *g_E_* and *g_I_*, are respectively the excitatory and inhibitory conductances, which increase by quantity *Q_E_* = 4 nS and *Q_I_* = 1.5 nS for each incoming spike. The increment of conductance is followed by an exponential decrease according to the equation:

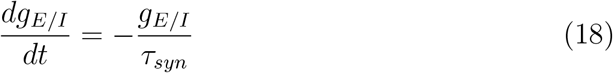

where *τ_syn_* = 5 ms. The spikes trains are generated by a Poissonian process with firing rate modulated by an Ornstein-Ulhenbeck stochastic process (Fourcaud & Brunel, 2002).

### A.4 I_M_ and I_h_ currents

The currents I_M_ and I_h_ are proposed as an example of currents which can be simulated using the CAdEx model. For the hyperpolarizing current I_h_, the parameters have been identified in (Banks et al., 1993) and for I_M_ the parameters have been identified in (Brown & Adams, 1980). The values of parameters for these currents are listed in Table 2.

**Table 2:** Parameters of the CAdEx to model neuron with I_M_ and I_h_ currents.

| | $C_m$ [pF] | $E_A$ [mV] | $E_L$ [mV] | $I_s$ [pA] | $V_A$ [mV] | $\Delta_A$ [mV] |
| --- | --- | --- | --- | --- | --- | --- |
| Neuron with $I_M$ | 200 | -90 | -60 | 350 | -35 | 4 |
| Neuron with $I_h$ | 200 | -43 | -60 | 100 | -75.7 | -5.7 |

Table 2: Parameters of the CAdEx to model neuron with $I_M$ and $I_h$ currents.
| | $V_R$ [mV] | $V_T$ [mV] | $\delta g_A$ [nS] | $\bar{g}_A$ [nS] | $g_L$ [nS] | $\tau_A$ [ms] |
| --- | --- | --- | --- | --- | --- | --- |
| Neuron with $I_M$ | -58 | -45 | 1 | 16 | 10 | 550 |
| Neuron with $I_h$ | -70 | -50 | 1.5 | 43 | 12 | 800 |

### A.5 Network simulations

The parameters of AdEx and CAdEx networks for which results are presented in Fig. 1 are shown in Tables 3, 4 and 5. The input to both networks was given by current noise injected to all neurons and had the form of an Ornstein-Uhlenbeck processes, with the same parameters: a variance, *σ* = 2.4 pA and a time constant, *τ* = 100 ms. The neurons were connected by conductance-based synapses (see Appendix A3).

**Table 3:** AdEx network parameters

|  |  | **AdEx** |  |
| --- | --- | --- | --- |
|  |  | Excitatory cells | Inhibitory cells |
| Population | $N$ | 800 | 200 |
| Capacitance | $C$ | 150 pF | |
| Leak conductance | $g_L$ | 10 nS | |
| Leak reversal potential | $E_L$ | -63 mV | -65 mV |
| Spike threshold | $V_T$ | -50 mV | |
| Spike initiation slope | $\Delta_T$ | 2 mV | 0.5 mV |
| Adaptation time constant | $\tau_w$ | 500 mV | n.a. |
| Post-spike adaptation increment | $b$ | 107 pA | n.a. |
| Subthreshold adaptation | $a$ | 0 nS | n.a. |
| Quantal synaptic conductance | $Q_{E/I}$ | 1.2 nS | 5 nS |
| Synaptic reversal potential | $E_{E/I}$ | 0 mV | -75 mV |

**Table 4:**
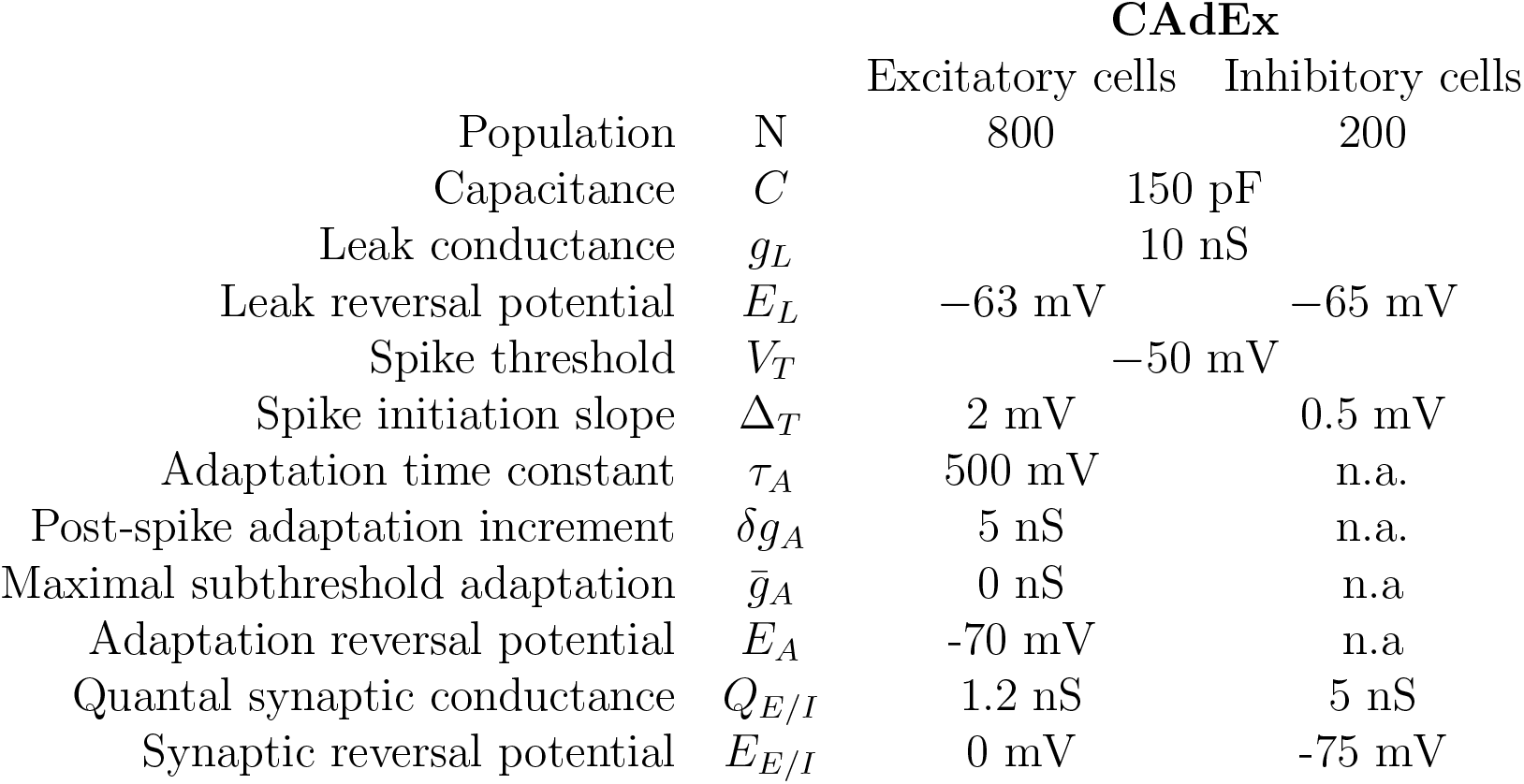
CAdEx network parameters

**Table 5:** Network connectivity

|  | Excitatory cells | Inhibitory cells |
| --- | --- | --- |
| Excitatory cells | 12% | 10% |
| Inhibitory cells | 10% | 12% |

### A.6 Unbounded hyperpolarization of the AdEx model

In the AdEx model the membrane voltage can be unbounded from below i.e. tends to minus infinity. This situation can occur if *a* is negative and *g_L_* < |*a*|, see Fig. A2. In this situation, the slope of the oblique asymptote of the *V* nullcline, which equals *g_L_*, is less steep than the slope of the adaptation nullcline. In the shaded area between nullclines in Fig. A2, the voltage tends towards minus infinity.

This divergence of the AdEx model to minus infinity is automatically fixed by the CAdEx model because the voltage is bounded to the reversal value of the adaptation current.

**Figure A2:**
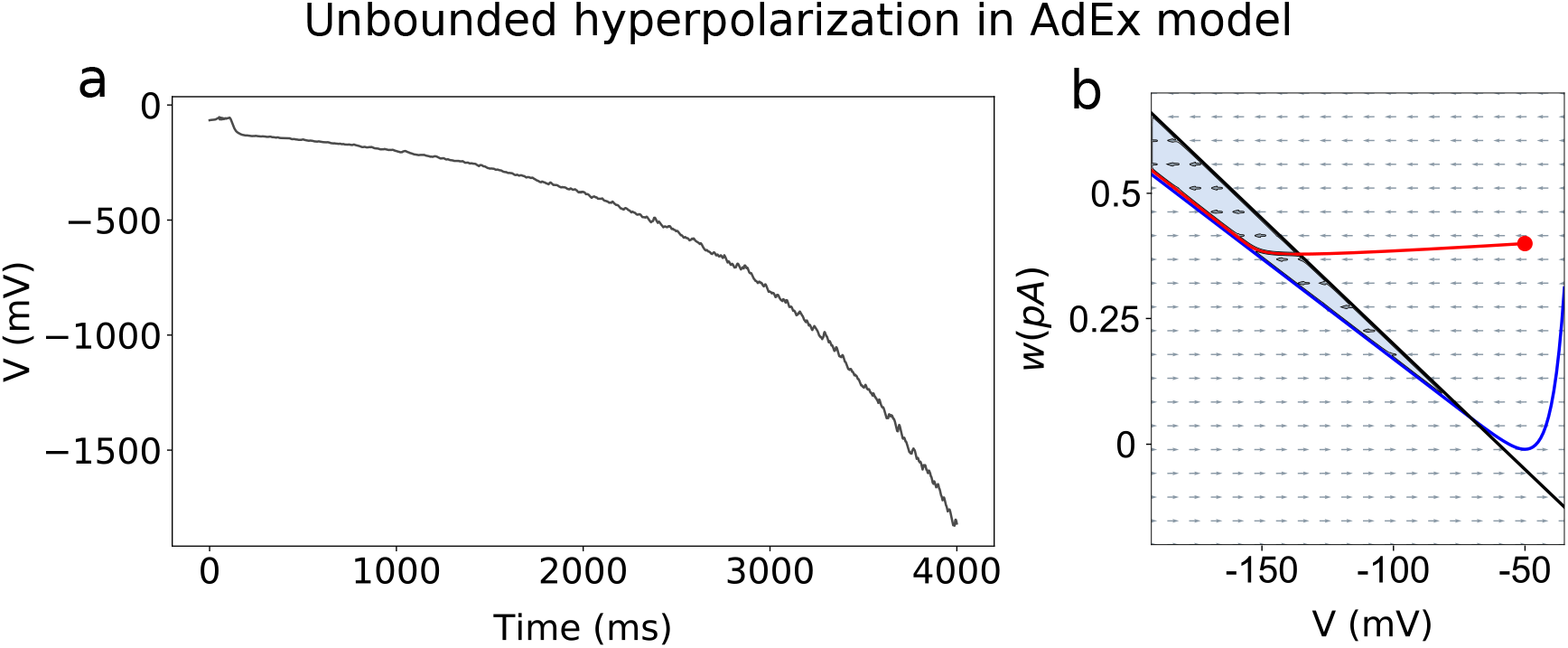
Unbounded hyperpolarization of the AdEx model. (a) Network simulation. The mean membrane voltage of excitatory AdEx neurons tends to minus infinity. The parameters of the network are the same as in Fig. 1b, see Table. 3, the only difference is the value of *a* parameter, here *a* = −15 nS (b) Corresponding phase plot. *Blue line* - voltage nullcline, *black line* - adaptation nullcline, *red line* - trajectory of the system.

### A.7 Mathematical overview of the AdEx model

The equations of the Adaptive Exponential Integrate and Fire (AdEx) Model are as follows:

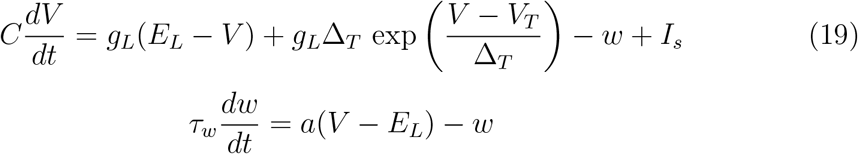

with the after-spike reset mechanism:

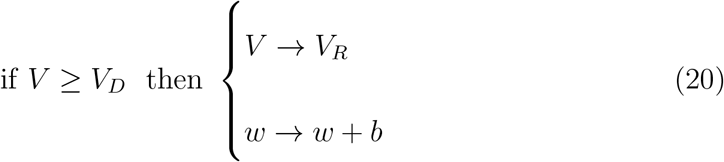

The nullclines of AdEx model are as follows:

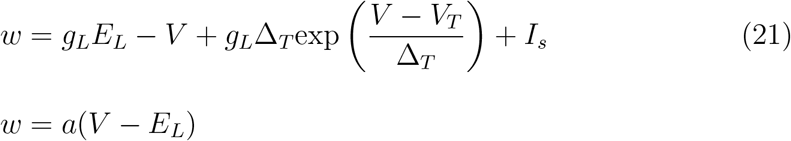

The Jacobian of the AdEx system around equilibrium *i*, located at (*V*^(*i*)^, *w*^(*i*)^) for an input current *I_s_*, has the form:

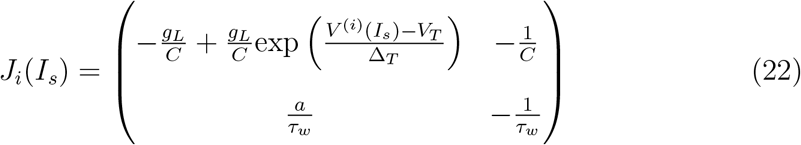

The trace 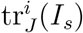 and the determinant 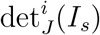 of the Jacobian are as follows:

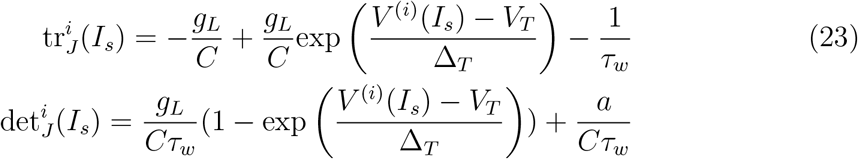

From above equations we get:

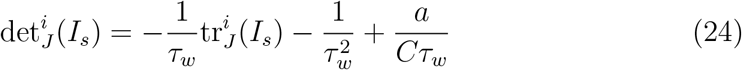

which tells us that the relationship between det_*J*_ and tr_*J*_ is linear in the AdEx model, i.e. the trajectories of the equilibria on the Poincarée diagram are straight lines.

## References

Allen Brain Institute. (2015). Allen cell types database. Retrieved from https://celltypes.brain-map.org/data

Banks, M. I., Pearce, R. A., & Smith, P. H. (1993). Hyperpolarization-activated cation current (ih) in neurons of the medial nucleus of the trapezoid body: voltage-clamp analysis and enhancement by norepinephrine and camp suggest a modulatory mechanism in the auditory brain stem. Journal of neurophysiology, 70(4), 1420–1432.

Benda, J., & Herz, A. V. M. (2003). A universal model for spike-frequency adaptation. Neural Computation, 15 (11), 2523–2564. doi: 10.1162/089976603322385063

Brette, R., & Gerstner, W. (2005). Adaptive exponential integrate-and-fire model as an effective description of neuronal activity. Journal of neurophysiology, 94(5), 3637–3642.

Brown, D., & Adams, P. (1980). Muscarinic suppression of a novel voltagesensitive k+ current in a vertebrate neurone. Nature, 283(5748), 673.

Clopath, C., Jolivet, R., Rauch, A., Lüscher, H.-R., & Gerstner, W. (2007). Predicting neuronal activity with simple models of the threshold type: Adaptive exponential integrate-and-fire model with two compartments. Neurocomputing, 70(10-12), 1668–1673.

Connor, J. A., & Stevens, C. F. (1971, Feb). Prediction of repetitive firing behaviour from voltage clamp data on an isolated neurone soma. J. Physiol. (Lond.), 213(1), 31–53.

Fitzhugh, R. (1961, Jul). Impulses and Physiological States in Theoretical Models of Nerve Membrane. Biophys. J., 1 (6), 445–466.

Fourcaud, N., & Brunel, N. (2002, Sep). Dynamics of the firing probability of noisy integrate-and-fire neurons. Neural Comput, 14 (9), 2057–2110.

Izhikevich, E. (2007). Dynamical systems in neuroscience: The geometry of excitability and bursting.

Izhikevich, E. M. (2003). Simple model of spiking neurons. IEEE Transactions on neural networks, 14 (6), 1569–1572.

Krinskii, V. I., & Kokoz, I. u. M. (1973). [Analysis of the equations of excitable membranes. I. Reduction of the Hodgkins-Huxley equations to a 2d order system]. Biofizika, 18(3), 506–511.

McCormick, D. A., & Contreras, D. (2001). On the cellular and network bases of epileptic seizures. Annu. Rev. Physiol., 63, 815–846.

Morris, C., & Lecar, H. (1981, Jul). Voltage oscillations in the barnacle giant muscle fiber. Biophys. J., 35(1), 193–213.

Naud, R., Marcille, N., Clopath, C., & Gerstner, W. (2008). Firing patterns in the adaptive exponential integrate-and-fire model. Biological cybernetics, 99 (4-5), 335.

Stimberg, M., Brette, R., & Goodman, D. F. (2019, aug). Brian 2, an intuitive and efficient neural simulator. eLife, 8, e47314. doi: 10.7554/eLife.47314

Touboul, J., & Brette, R. (2008, Nov). Dynamics and bifurcations of the adaptive exponential integrate-and-fire model. Biol Cybern, 99(4-5), 319–334.

Treves, A. (1993). Mean-field analysis of neuronal spike dynamics. Network: Com-putation in Neural Systems, 4 (3), 259–284. doi: 10.1088/0954-898X_4_3_002

